# Aer is a bidirectional redox sensor mediating negative chemotaxis to antibiotic-induced ROS in *E. coli*

**DOI:** 10.1101/2025.11.30.691477

**Authors:** Nabin Bhattarai, Shelley Payne, Rasika M. Harshey

## Abstract

The Aer chemoreceptor in *E. coli* is known to perform oxygen- or aero-taxis by sensing metabolic flux through ETC via an FAD cofactor bound to its PAS domain. We show in this study that Aer also senses oxidative stress and performs FAD-dependent negative chemotaxis in response to the known ROS H_2_O_2_. The ability of Aer to detect both oxidizing and reducing intracellular environments redefines its functional range and establishes Aer as a bidirectional redox sensor. We show in addition that the ability of bacterial swarms to move away from bactericidal antibiotics is Aer-dependent and is abolished by expression of the catalase–peroxidase enzyme KatG, which scavenges intracellular ROS. Our study thus provides independent behavioral evidence that certain antibiotics generate intracellular ROS, while also offering a sensitive assay for detecting these reactive species.

**Importance:** Motile bacteria rely on aerotaxis to seek environments that maximize energy production. We show that in *E. coli*, Aer mediates not only positive chemotaxis toward favorable redox conditions but also negative chemotaxis in response to oxidative stress. The ability of Aer to detect both oxidizing and reducing cellular environments reveals an unexpected sensory versatility, shared to varying degrees by other *E. coli* chemoreceptors. We show that *E. coli* uses Aer to actively move away from ROS-generating antibiotics, revealing a previously unrecognized behavioral mechanism for bacterial survival under oxidative stress.

## Introduction

*E. coli* employs chemotaxis to sense and navigate its surroundings with the help of 4-6 peritrichous flagella, driven at their base by bidirectional rotary motor powered by PMF (proton motive force) (1–3). The chemotaxis machinery controls the direction of flagellar rotation, allowing the bacteria to move towards favorable environments or away from unfavorable ones. Chemoreceptors, also called MCPs (methyl-accepting chemotaxis proteins), sense specific ligands in the periplasm and convey this information to a linked cytoplasmic kinase CheA (4). Attractants and repellents change MCP conformation to either inhibit or activate CheA autophosphorylation (CheA-OFF or ON, respectively). CheA-ON phosphorylates two response regulators – CheY and CheB. CheY∼P interacts with the bidirectional flagellar motor to change its default counterclockwise (CCW) rotation direction to clockwise (CW) (1). Phosphatase CheZ dephosphorylates CheY∼P to terminate the activation signal, restoring CCW rotation. The directionality of motor rotation therefore reports on CheA activity. Every signaling event is followed by an adaptation event that resets the pre-signaling conformation of the MCPs. This is achieved by CheB and CheR. CheB∼P is a methylesterase, which preferentially interacts with the CheA-ON state of receptors and shifts them toward the OFF state, while CheR is a methyltransferase that preferentially interacts with CheA-OFF state of the receptors and methylates E residues in the MCPs to return them to a CheA-ON state (4).

*E. coli* possesses a repertoire of five dimeric chemoreceptors- Aer, Tsr, Tar, Tap and Trg - that sense different ligands. Aer responds to changes in oxygen concentration, mediating aerotaxis not by direct interaction with O_2_, but by monitoring the redox state of the electron transport chain (ETC) (5–9). Unlike the other four chemoreceptors, Aer is not an MCP, in that it lacks the E residues important for methylation-based adaptation (10, 11). Also unique is its sensor PAS (Per-Arnt-Sim) domain in the cytoplasm (Fig. 1). PAS binds cofactor FAD (flavin adenine dinucleotide) (10, 12). The HAMP domain plays a crucial role in stabilizing the PAS domain, and the interaction between these domains is vital for the optimal functionality of Aer (13, 14). Redox-dependent conformational changes in the PAS domain alter Aer’s HAMP and kinase interaction domains, modulating CheA activity and thus rotor bias. The differing redox states of FAD, including FAD^2+^ and FADH, when bound to Aer, have been reported to control its signaling output *in vitro* (15, 16). As measured by phosphorylation of CheY, FAD^2+^ activates, and FADH (anionic semiquinone or ASQ) inactivates CheA; Aer-FADH_2_ (hydroquinone or HQ) has not been observed *in vitro* (15, 17).

**Fig. 1.**
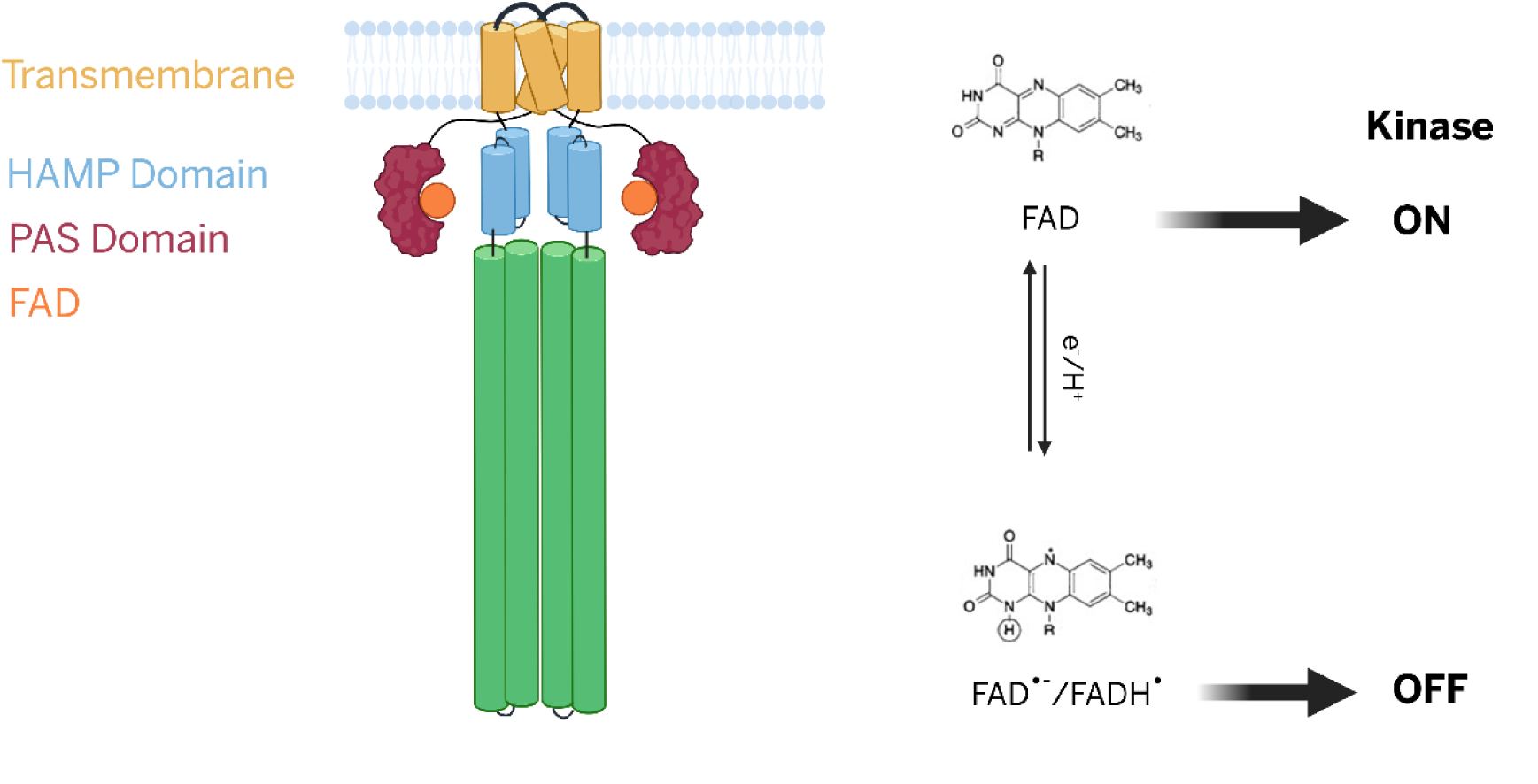
Model of Aer-mediated redox signaling and kinase regulation. Schematic representation of the Aer dimer showing the organization of signaling domains: HAMP (blue), PAS (burgundy), bound FAD cofactor (orange). The redox state of FAD bound within the PAS domain determines Aer signaling output. Fully oxidized FAD maintains the kinase in the ON state. One-electron reduction forms the anionic or neutral semiquinone (FAD^•^⁻/FADH^•^), switching the kinase OFF.

The present study was motivated by our prior observation that inclusion of antibiotics kanamycin or ciprofloxacin in the swarm medium, induced a CheY-dependent avoidance or repellent response within a *Serratia marcescens* swarm, an effect not attributable to the antibiotics acting as ligands *per se* (18). Bactericidal antibiotics are known to generate reactive oxygen species (ROS) as secondary consequences of metabolic perturbations involving the TCA cycle and electron transport chain, through damage to iron–sulfur clusters in proteins, and via stimulation of the Fenton reaction (19–21). Notably, *Helicobacter pylori* has been shown to swim away from H₂O₂, a known source of ROS (22). Because Aer regulates CheA activity in response to cellular redox changes, we first examined the rotational behavior of individual flagellar motors to monitor the Aer response to ROS and subsequently assessed its response to antibiotics. We find that H₂O₂, a canonical ROS, elicits an immediate chemorepellent response, whereas ciprofloxacin and kanamycin trigger a slightly delayed response. Aer was identified as the primary mediator of these effects, with FAD being essential for its function. These findings broaden the known repertoire of Aer activities and provide new assays for monitoring ROS-dependent signaling.

### *E. coli* responds to H_2_O_2_ as a chemorepellent

Since the direction of motor rotation reflects CheA activity, we used the bead assay for motor rotation (see Methods), to examine the effect of ROS. The *E. coli* strain MG1655 (WT) and its derivatives were used in most assays. Under basal conditions, WT motors exhibited the expected bias, rotating predominantly in the CCW direction with frequent CW reversals (39.07±1.75 reversals per minute or rpm) (23) (Fig. 2A, left). Motor responses to hydrogen peroxide (H₂O₂) were monitored at 1 mM, a concentration previously used in studies reporting antibiotic-induced ROS production (19) and repellent responses to H₂O₂ in *H. pylori* (22). Upon H₂O₂ addition, motor reversals increased significantly (64.53 ± 3.93 rpm), indicative of a repellent response (Fig. 2A, right and Fig. S1). The motors could not be followed beyond one minute because the bead invariably detached from the filament stub. Titration experiments showed that as little as 10 µM H₂O₂ was sufficient to elicit a significant repellent response (Fig. S1B).

**Fig. 2.**
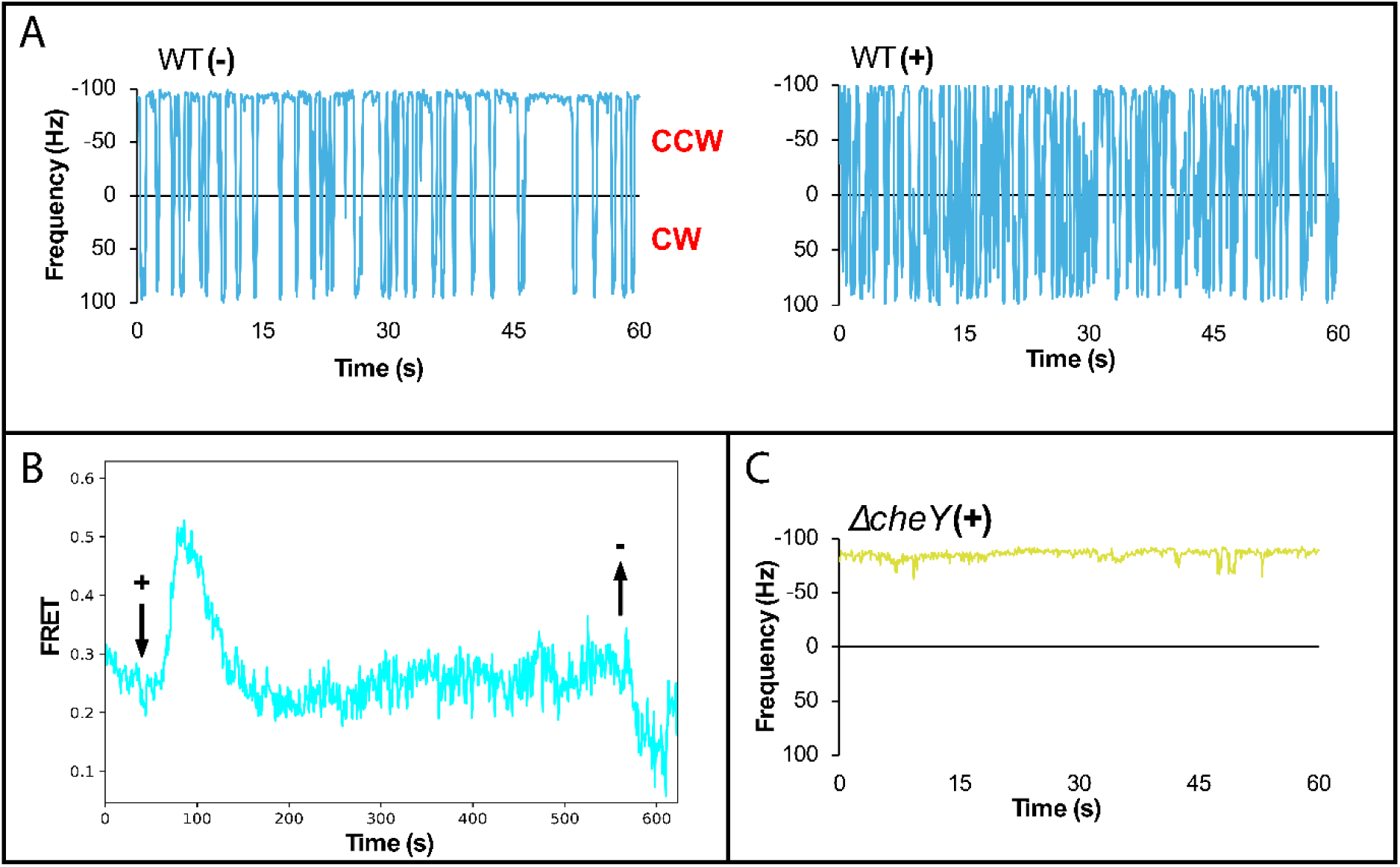
H₂O₂ elicits a repellent response in *E. coli* as measured by two assays. (A) Representative motor trace of WT (NBN58) *E. coli* before (-) and immediately after (+) addition of 1 mM H₂O₂. See Fig. S1 for a summary of the cumulative data. (B) FRET measurements in VS115 showing population level CheA activity (see Methods). The addition and removal of 1 mM H₂O₂ are indicated by downward (+) and upward arrows (-). The trace represents the average of three independent experiments. (C) As in A except with Δ*cheY*.

A complementary assay based on Förster resonance energy transfer (FRET), which monitors CheA activity through the interaction of CheY and CheZ fused to donor and acceptor fluorophores (see Methods), reproduced the motor response observed in the bead assay: CheA activity increased immediately upon H₂O₂ addition (Fig. 2B). In addition, this assay revealed that CheA activity returned to baseline within approximately one minute, consistent with adaptation to the chemosensory response.

To determine whether the response to H₂O₂ required chemotaxis signaling, we repeated the experiment using a Δ*cheY* mutant, which exhibits an extreme CCW bias due to the absence of the CW signal CheY∼P (Fig. 2C). In this strain, H₂O₂ failed to elicit any motor response, confirming that the repellent response to H₂O₂ observed in the WT is mediated through the canonical chemosensory pathway. A summary of motor traces for WT and the *cheY* mutant is shown in Fig. S1A.

### Response to peroxide is via Aer

Aer is thought to monitor the cellular redox state through its FAD cofactor, with the fully oxidized form activating CheA *in vitro* (15). To test whether Aer plays a similar role *in vivo*, we examined the response of a Δ*aer* strain to H₂O₂. The basal motor bias of Δ*aer* was comparable to that of the WT [Fig. 3A, top left (compare to Fig. 2A left) and Fig. S2A]. While WT cells showed a robust repellent response to H₂O₂ (Fig. 3A, bottom left), this response was abolished in the *aer* mutant (Fig. 3A, top right). Interestingly, instead of returning to the basal bias, the mutant now exhibited a clear attractant response.

**Fig. 3.**
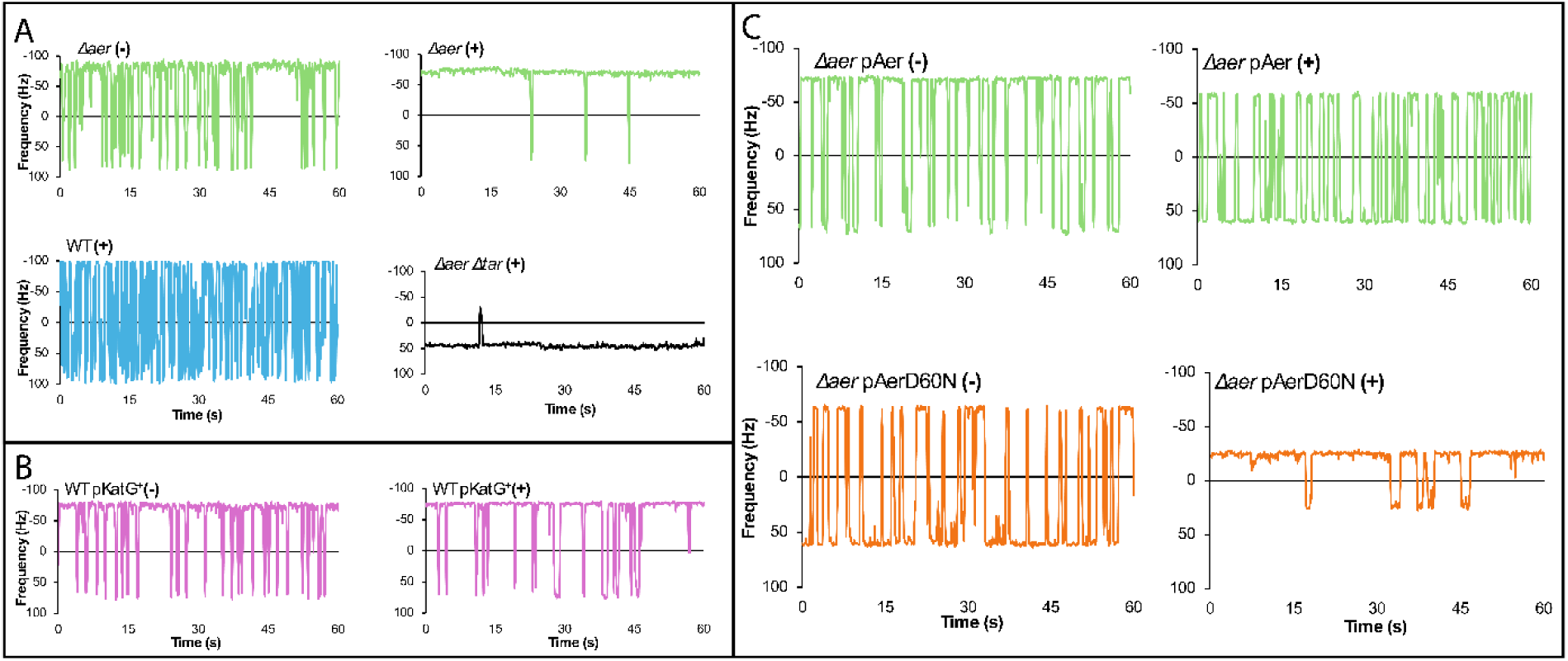
Chemorepellent response to peroxide is via Aer. (A) Representative motor traces of WT *E. coli* and its Δ*aer* (NBN30) and Δ*aer* Δ*tar* (NBN130) variants, with (+) and without (-) H₂O₂ addition. Experimental protocol as in Fig. 2A. (B) Response to H₂O₂ of WT harboring p*katG* induced (+) with IPTG. (C) Comparison of H₂O₂ responses of pAer and pAerD60N introduced into Δ*aer* and induced with IPTG.

Because H₂O₂ is a weak acid (pH 6.2 at 1 mM), we considered whether this attractant response might arise from pH sensing by the MCPs Tar and Tsr, known to exhibit opposing responses to pH: Tar senses low pH as an attractant, whereas Tsr mediates a repellent response (24). To test whether Tar contributed to the attractant response in the Δ*aer* mutant, we constructed an *aer tar* double mutant. As expected, this strain displayed a repellent response to low pH, attributable to Tsr (Fig. 3A, bottom right; Fig. S2A).

To determine whether the WT response to H₂O₂ reflected a combined input from Aer responding to ROS and Tsr responding to pH, we conducted two additional tests. First, we introduced into the WT strain a plasmid expressing the catalase–peroxidase KatG, which degrades intracellular H₂O₂(25). Induction of *katG* with IPTG eliminated the repellent response to H₂O₂ (Fig. 3B; Fig. S2B), indicating that the WT repellent response is due to ROS rather than to pH (note that the motor now shows a more CCW behavior, consistent with Tar sensing low pH as an attractant). Second, because Aer depends on FAD binding to its PAS domain for energy taxis (5), we examined the role of this cofactor using the AerD60N mutant, which is unable to bind FAD (10) and is defective in aerotaxis (11). The *aer* deletion strain complemented with WT Aer (pAer) exhibited the expected repellent response to H₂O₂ (Fig. 3C, top; compare left and right traces). In contrast, the FAD-binding–deficient mutant (pAerD60N) displayed the attractant response characteristic of Tar (Fig. 3C, right traces). Notably, this mutant also exhibited an intrinsic CW bias (Fig. 3C, bottom left), reminiscent of CW-biased, aerotaxis-defective HAMP-domain mutants of Aer (26). These data are summarized in Fig. S2C. We note that Δ*aer* or AerD60N cells exposed to H₂O₂ consistently exhibited lower motor speeds (Fig. S3A,B) (see Discussion).

Together, these findings demonstrate that FAD binding to Aer is essential for eliciting a repellent response to H₂O₂ and establish Aer as an ROS sensor in addition to its known role as an O₂ sensor.

### *E. coli* shows a ROS-dependent repellent response to kanamycin

Having established that *E. coli* exhibits a chemosensory response to ROS via Aer, we next tested whether *E. coli* swarms display the avoidance response to sublethal levels of kanamycin and ciprofloxacin previously observed in *S. marcescens* swarms (18). Using defocused epifluorescence video imaging, in which the z-position of GFP-labeled cells within the swarm colony is inferred from the diameter of their diffraction ring (18), we compared the distribution of GFP-labeled WT cells and their Δ*aer* variant grown on swarm medium containing sublethal kanamycin (3 µg/ml) (see Methods). The average colony height at the observed region in the swarm was approximately 150 µm. When viewed from above, GFP-cells located near the agar surface appear in focus, whereas cells positioned higher in the colony display a fluorescent diffraction ring whose diameter increases with the distance from the bottom surface. WT *E. coli* cells preferentially localized away from the agar surface, whereas the *aer* mutant cells were uniformly distributed throughout the colony (Fig. 4A). These results with *E. coli* mirror those reported for *S. marcescens* swarms (18), identifying in addition that Aer is responsible for the avoidance response.

**Fig. 4.**
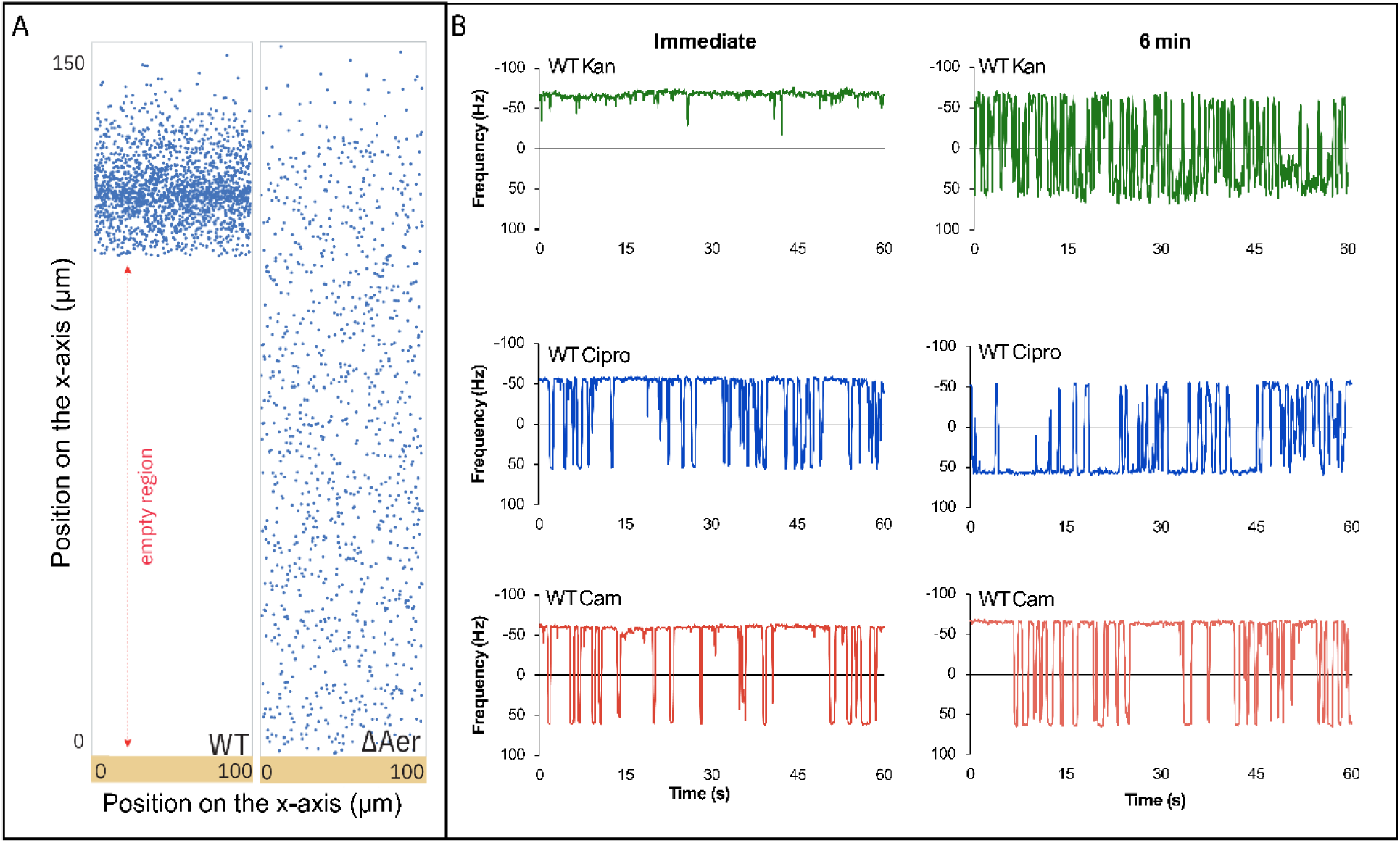
*E. coli* cells show an Aer-dependent avoidance response to kanamycin within a swarm. (A) Z-axis tracking of GFP-labeled cells in WT (NBN66) and *Δaer* (NBN67) swarms grown on medium containing 3 µg/mL kanamycin as described under Methods. The Y-axis represents the vertical position within the colony (total height ≈150 µm), while the X-axis indicates the horizontal position of each tracked cell. ‘Empty region’ refers to absence of GFP-labeled cells. Data were collected from >10 experiments ending in ∼1000 points each. Statistical significance was calculated against the no-treatment control using paired students t-test (\*\**p<0.01*; \*\*\**p<0.001*). (B) Representative motor traces of WT *E. coli* treated with kanamycin (10 µg/ml), ciprofloxacin (1 µg/ml), and chloramphenicol (15 µg/ml). Motors were tracked continuously for 10 minutes after addition of the antibiotic.

Since bactericidal antibiotics are known to induce ROS in bacteria, we next monitored the motor responses of *E. coli* cells to sublethal concentrations of kanamycin and ciprofloxacin - antibiotics reported to generate ROS - and to chloramphenicol, a bacteriostatic antibiotic that reportedly does not (19). Because antibiotics generate ROS as a secondary effect of their primary targets, we did not expect an immediate response as seen with exogenous H₂O₂. Accordingly, motor rotation was tracked for 10 min following antibiotic addition in WT cells. Kanamycin elicited a biphasic response: an immediate attractant phase (Fig. 4B, top panel, left), followed by a delayed repellent phase beginning at approximately 6 min (Fig. 4B, top panel, right). Ciprofloxacin induced a similar delayed repellent response but lacked the initial attractant phase (Fig. 4B, middle panel). In contrast, chloramphenicol elicited no detectable response within this time window (Fig. 4B, bottom panel).

The transient attractant response to kanamycin could reflect an early perturbation of cellular respiration, in which case reduced oxygen consumption can decrease ETC activity, leading to accumulation of reduced electron carriers (NADH, FADH₂) and a more reduced intracellular redox state, favoring in turn a reduction of the FAD cofactor in Aer (26). This phase is subsequently followed by increased oxidative activity and ROS accumulation (27), producing oxidized FAD and a corresponding repellent response as early as 6 minutes after antibiotic exposure.

To determine whether the repellent response to kanamycin is Aer- and ROS-dependent, we repeated the experiment using a Δ*aer* mutant, as well as in WT cells expressing the catalase–peroxidase KatG. The *aer* mutant failed to exhibit either the early attractant or late repellent response (Fig. 5A, top). In WT cells expressing KatG, the delayed repellent phase was abolished, whereas the initial attractant phase persisted (Fig. 5B, bottom). These results confirm that the antibiotic-induced chemorepellent response is mediated through Aer and depends on ROS signaling. A summary of the data presented in Fig. 5 is shown in Fig. S4.

**Fig. 5.**
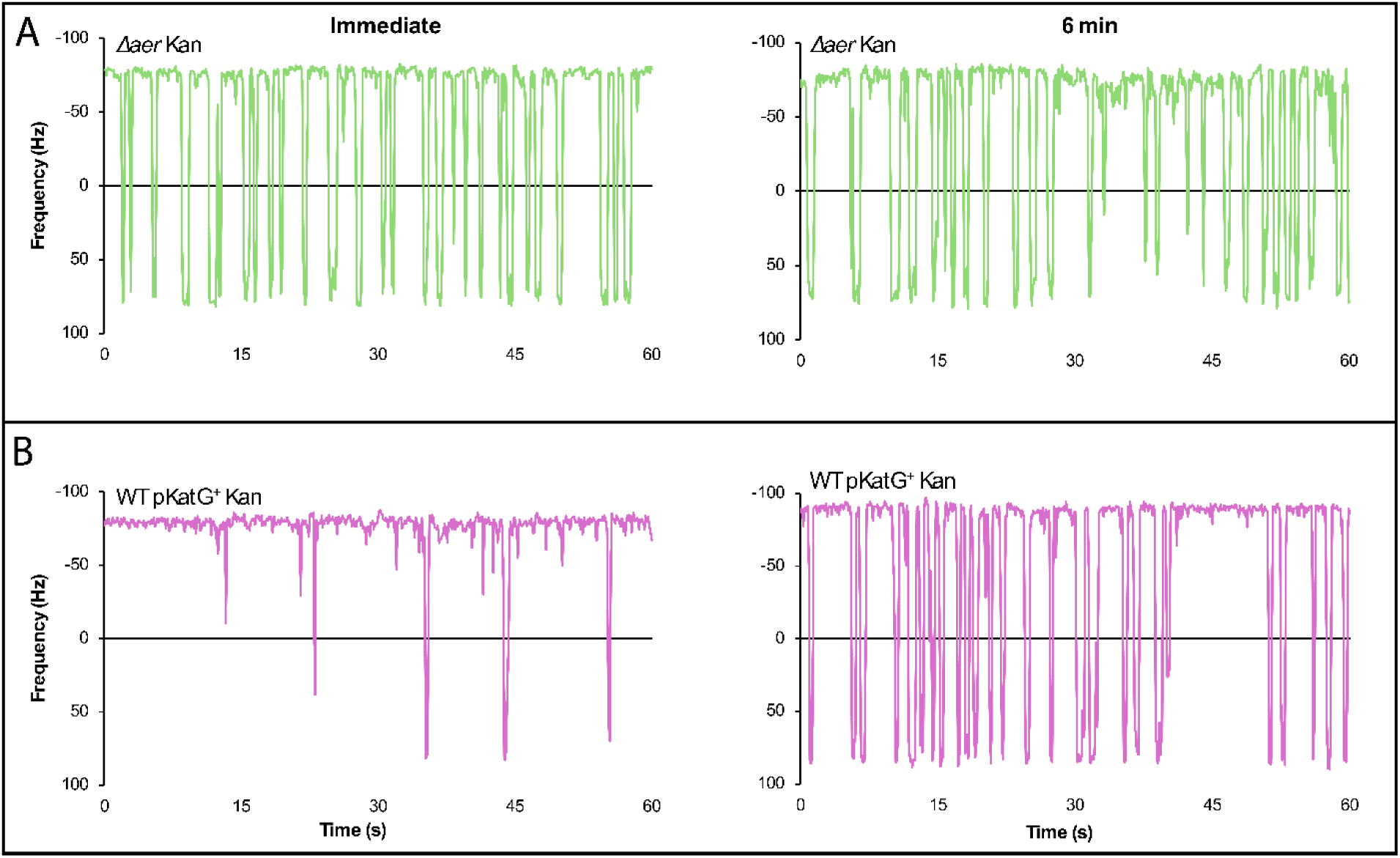
Repellent antibiotic response is Aer and ROS-dependent. Representative motor traces of (A) Δ*aer* (NBN 30) and (B) WT harboring p*katG* (NBN 33) treated with kanamycin (10 µg/ml). Motors were tracked continuously for 10 minutes after addition of the antibiotic. Other labels as in Fig. 4.

## Discussion

The results presented in this study expand the functional repertoire of the chemoreceptor Aer. Using two complementary assays— motor behavior and FRET—we show that Aer, previously known to mediate positive chemotaxis toward O₂ in *E. coli*, also mediates negative chemotaxis to ROS (Figs. 2, 3). These assays provide a sensitive and convenient means to monitor intracellular ROS. We further used Aer’s ROS-sensing ability to support the growing consensus that bactericidal antibiotics, in addition to their primary targets, kill through ROS generated in downstream metabolic reactions (21).

Aer is distinct among *E. coli*’s five chemoreceptors: it is expressed at only ∼10% of the level of the major receptors Tsr and Tar (whose estimated copy number ranges from 10,000 to 30,000 per cell (28)), and lacks their methylation-based adaptation mechanism (5, 10). Yet Aer adapts to O₂ stimuli in soft agar assays (11). Using FRET, we now show that Aer can also adapt to ROS signals (Fig. 2B). The observed repellent response to ROS is consistent with *in vitro* findings showing that Aer bound to the reduced FADH form signals an attractant response, whereas the oxidized FAD form signals repulsion (15). *In vivo* exposure to H₂O₂ is expected to shift the FAD pool toward the oxidized state, consistent with our behavioral data. Present at an estimated 1000-3000 copies per cell, Aer may sequester a substantial portion of cellular FAD. Given our observation of consistently lower motor speeds in Δ*aer* cells exposed to H₂O₂ (Fig. S3A, B), we speculate that in the absence of Aer, elevated cytoplasmic FAD levels could compete with endogenous flavoproteins or quinones for reducing equivalents, effectively diverting electrons from the ETC and lowering PMF, which powers the motor. Figure 6 summarizes our understanding of Aer as advanced in this study.

**Fig. 6.**
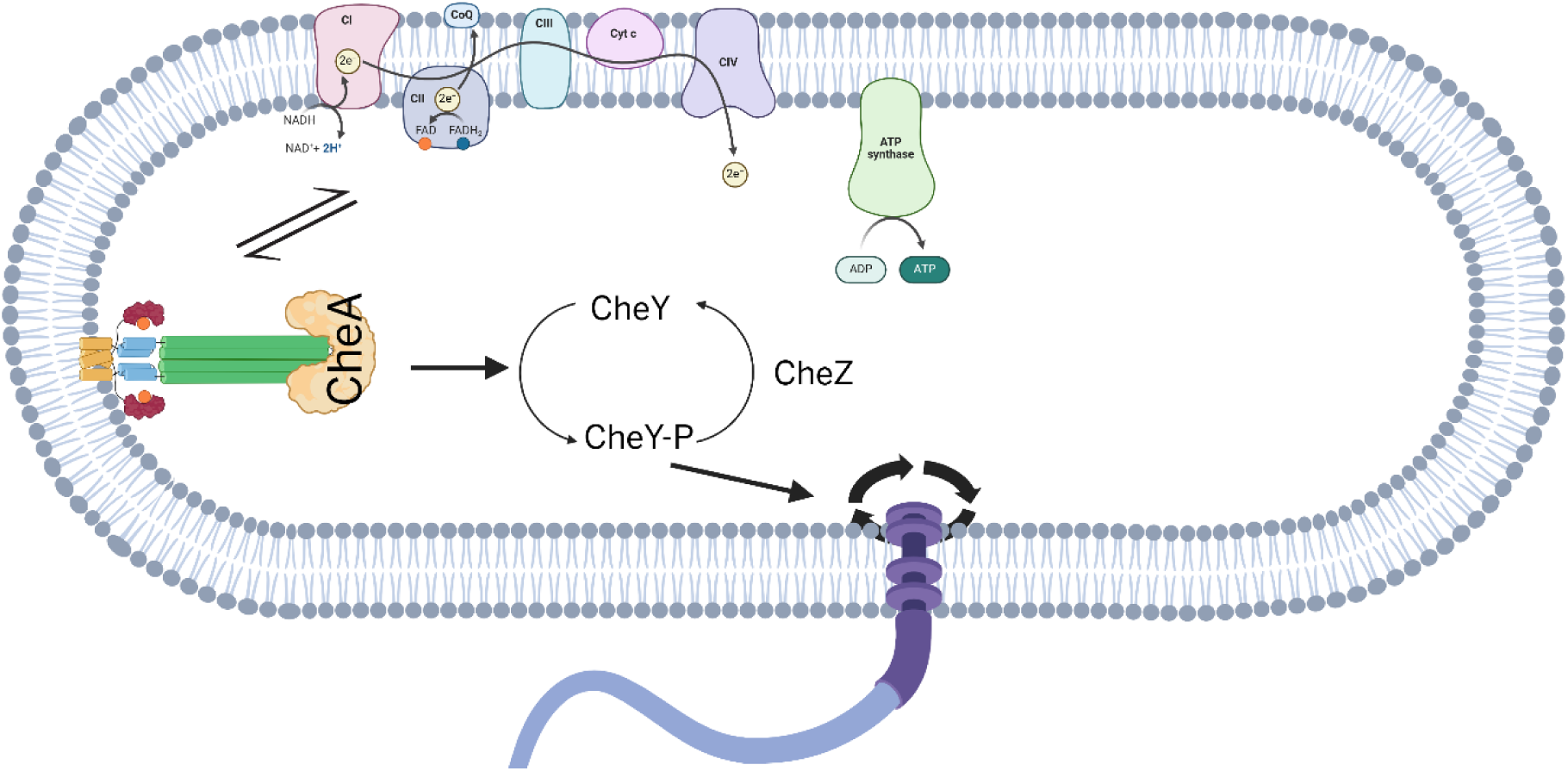
Summary. Aer-bound FAD/FADH reflects global electron transport chain activity by equilibrating with the cytoplasmic redox state in *E. coli*, mediating both positive chemotaxis toward O₂ and negative chemotaxis to ROS.

We also replicated in *E. coli* the avoidance responses to kanamycin and ciprofloxacin previously observed in *S. marcescens* swarms (18), and demonstrated that the response to kanamycin is Aer-dependent (Fig. 4A). Although the role of ROS in antibiotic lethality was once debated (29), accumulating evidence supports the idea that accelerated metabolism and consequent ROS generation are central to the action of bactericidal antibiotics (21). ROS production following antibiotic treatment has typically been monitored using genetically encoded sensors (30), oxidation-sensitive dyes (19, 31), or oxygen consumption assays (27), which detect ROS over tens of minutes. In contrast, Aer-mediated CheA activation by H₂O₂ occurred immediately in both bead and FRET assays (Fig. 2), and responses to sub-lethal amounts of bactericidal antibiotics were observed within minutes (Fig. 4B), providing a more rapid and sensitive measure of ROS generation. Notably, no ROS response was detected to the bacteriostatic antibiotic chloramphenicol within the 10-minute observation window (Fig. 4B), consistent with its distinct mode of action.

The biphasic motor response to kanamycin—an initial attractant phase followed by a delayed repellent phase, with only the latter abolished by KatG expression (Fig. 5B) - highlights the dynamic nature of Aer-mediated signaling. These results suggest that kanamycin likely initially suppresses electron transport, leading to accumulation of reduced equivalents and a more reduced Aer-bound FAD, followed by increased O₂ consumption and ROS generation that reoxidizes FAD. The differences in the motor responses of the three structurally different antibiotics tested reveal differences in their effects on cellular redox.

The role of Aer in ROS sensing parallels that of *H. pylori*’s TlpD, a cytoplasmic chemoreceptor that mediates repulsion from oxidative stress to help the bacterium evade host-derived ROS (22). TlpD has no identifiable domains that would bind FAD/FADH, but possesses a CZB domain at its C terminus, which binds zinc via histidine and cysteine residues (32). Other ROS-sensing proteins in both prokaryotes and eukaryotes use cysteine as a way to sense redox status (33), so this mechanism is speculated to operate in TlpD as well (32).Together, these findings provide insight into how bacteria might evade ROS in their natural environments, whether host-derived or antibiotic-generated. By actively moving away, bacteria may reduce local drug exposure and oxidative damage, adding a behavioral dimension to antibiotic resistance strategies (Fig. 4A). Targeting chemotaxis pathways—or Aer signaling specifically—could therefore enhance bacterial susceptibility to ROS-generating antibiotics and limit their capacity to evade treatment.

## Acknowledgments

We thank Jeremy Moore and Thierry Emonet at Yale University for running the FRET experiment in Figure 2B and for discussion, Ajesh Jose in Avraham Be’er’s lab at the Ben-Gurion University of the Negev, Israel, for help with the experiment in Figure 4A, and S. Bhattacharyya, Y. S. Hwang, and N. Wadhwa for discussion. Figures 1 and 6 were generated using BioRender. This work was supported by NIH grant GM118085 to RMH.

## Methods

### Strains and Growth Conditions

All strains and plasmids used in this study are listed in Tables S1 and S2. *E. coli* cultures were grown in Lennox Broth (LB; 20 g/L, Fisher BioReagents). Antibiotics were added at the following concentrations for strain selection: ampicillin (100 µg/mL), chloramphenicol (20 µg/mL), and kanamycin (50 µg/mL). For plasmid induction, 50 µM IPTG (isopropyl-β-D-thiogalactopyranoside) or 0.2% L-arabinose was used, unless otherwise as indicated. Cell growth was monitored by measuring optical density at 600 nm (OD₆₀₀) using a spectrophotometer.

All FRET experiments were performed with RP437, a derivative of *E. coli* K-12 harboring CheYZ and *fliC* mutations and transformed with two plasmids: pSJAB106 from which CheZ-YFP and CheY-mRFP1 are expressed in tandem on IPTG inducible promoter, and pZR1 from which ‘sticky’ FliC^∗^ is expressed on a sodium-salicylate (NaSal) inducible promoter. Cells were grown overnight in TB broth (1% bacto-tryptone, 0.5% NaCl), then diluted 1:100 into 10mL of fresh TB supplemented with 50 μM IPTG to induce the FRET pair, and 3µM NaSal to induce sticky FliC^∗^ for cell adhesion to glass coverslips, with 100µg/mL ampicillin and 34µg/mL chloramphenicol for plasmid retention.

### Mutagenesis and Plasmid Constructions

The WT parent strains of *E. coli* used were MG1655. Mutant strains were generated by inserting a kanamycin resistance (KAN) cassette into the target gene using the Keio collection (34). Mutations were transferred to fresh strain backgrounds by P1 Cm transduction, and KAN cassettes were excised via FLP recombinase expression from pCP20 (35). All resulting strains were verified by DNA sequencing.

To construct plasmids of interest, specific genes were amplified from the wild-type (WT) strain and cloned into pBAD33 via Gibson Assembly (36). Site-directed mutagenesis of *aer* was achieved using primers containing base substitutions to amplify the entire pNBN4 plasmid. For inducible expression of KatG, we used the ASKA library plasmid (ID: JW3914) containing the corresponding open reading frame (ORF) on pCA24N (37).

### Bead Assay

Bead assays were performed as previously described (23, 38). *E. coli* Δ*fliC* strains were complemented with plasmid pFD313 expressing sticky FliC^∗^ under a constitutive promoter (39). Flagella were sheared by passing cells through two syringes connected by a 7-inch polyethylene capillary. A 40 µL cell suspension was applied to a poly-L-lysine–coated coverslip attached to the glass slide with double sided tape, incubated for 10 min, and washed with motility buffer to remove unattached cells (40). Subsequently, 40 µL of a 1:50 dilution of polystyrene beads was added, incubated for 10 min, and excess beads were washed away. Bead rotation was recorded at 1,000 frames/s using a CCD camera (ICL-B0620M-KC0; Imperx, Boca Raton, FL). Videos were analyzed with custom LabVIEW 2012 software as previously described (38).

### Ligand Exposure and Chemotactic Response Recording

Once a properly attached bead on a flagellar stub was identified within the microscope’s field of view, a 60-second baseline recording was obtained. Ligands were then introduced through the top of the chamber and gently drawn out from the bottom using a Kimwipe to maintain smooth flow. Post-exposure recordings were collected for 60 seconds for peroxide treatments and up to 10 minutes for antibiotic treatments, or until the bead detached from the flagellum. The concentrations of the antibiotics tested were 10 µg/mL kanamycin, 1 µg/mL ciprofloxacin, and 15 µg/mL chloramphenicol. For hydrogen peroxide, the concentrations tested were 1 mM, 100 µM, 50 µM, and 10 µM.

### Data Acquisition and Analysis

Bead assay data included both rotation frequency and direction for individual motors. Counterclockwise (CCW) rotations were assigned negative values, while clockwise (CW) rotations were positive. CW bias, defined as the fraction of time a motor rotated CW within a given interval, was calculated as the ratio of positive to total data points in that interval (41, 42).

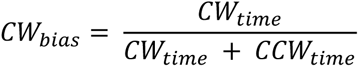

Batch processing was performed using a custom Python script (all the python code for this work is available at [https://github.com/nbnnpl01/CW-bias-calculation.git]). For long recordings, CW bias was smoothed using a 30 s running average. Statistical significance was determined using Student’s *t*-test comparing each sample to its corresponding WT control.

### Three-Dimensional Tracking of *E. coli* Cells within Swarming Colonies

Cells were grown on swarm plates containing 2.5 g/L LB, 0.45% agar, and 0.5% glucose, supplemented with 3µg/mL kanamycin. GFP-labeled cells (pJP10120) were mixed 1:100 with unlabeled cells prior to inoculation. A 20 µL inoculum was placed at the center of the plate, and colonies were grown at 30°C and 95% relative humidity for 9 h. Wild-type and Δ*aer* swarms of *E. coli* were grown and monitored separately but under identical conditions.

Swarm colonies (∼4 cm diameter) were harvested, and imaging was performed in the colony interior, ∼0.5 cm from the colony edge at a height of ∼150 µm. Cell Z-positions were determined using an off-focus diffraction ring method (18), where the ring diameter correlates with cell height (z-resolution ∼0.5 µm). Data were collected from multiple experiments (≈1,000 points per dataset) using a Zeiss Axio Imager Z2 microscope with a 63× objective (field of view = 100 µm × 100 µm) and a Neo Andor camera. Filter set 46 (YFP) was used instead of set 38 (GFP) to avoid light-induced pausing.

### *In Vivo* Single-Cell FRET Microscopy

Single-cell FRET imaging and sample preparation were performed as described previously (43). Cells were grown and washed as outlined above. Imaging was conducted using a Nikon Eclipse Ti-E inverted microscope with a 60× oil immersion TIRF objective (CFI Apo TIRF 60× Oil, Nikon). Yellow fluorescent proteins were illuminated with a SOLA SE LED system (Lumencore) through excitation filters (59026x, Chroma; FF01-500/24-25, Semrock) and a dichroic mirror (FF520-Di02, Semrock). Emission signals were split into donor and acceptor channels using an OptoSplit II system (Cairn) and collected through emission filters (FF01-542/27 and FF02-641/75; Semrock) on an ORCA-Flash 4.0 V2 camera (Hamamatsu). mRFP1 was imaged similarly, except with excitation filter FF01-575/05-25 and dichroic mirror FF593-Di03-25x36. All images were acquired with 50 ms exposure time.

### FRET Data Analysis

Single-cell fluorescence time-series were analyzed as previously described (43–45). Cells were segmented, and donor (D) and acceptor (A) signals were extracted using in-house software. Photobleaching was corrected by fitting donor and acceptor traces to bi-exponential decay functions.

To calculate FRET from fluorescence time-series, we used the E-FRET method.(44) The E-FRET index is given by

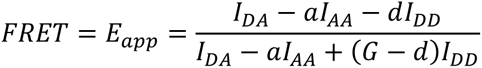

where 𝐼_𝐷𝐴_ is FRET-acceptor emission intensity from donor excitation, 𝐼_𝐴𝐴_ is acceptor emission with acceptor excitation, 𝐼_𝐷𝐷_ is donor emission from donor excitation, with 𝑎, 𝐺, 𝑑 being optical constants that depend on the FRET pair and optical setup which were determined by an independent experiment with strains that express only CheY-mRFP or only CheZ-mYFP.

FRET time-series were normalized for each cell using the minimum and maximum FRET values measured during saturating stimuli at the start and end of experiments to estimate kinase activity.

